# Insulin-like signalling influences the coordination of *Drosophila* larval hemocyte number with body size

**DOI:** 10.1101/2020.02.05.935098

**Authors:** Daniel Bakopoulos, Lauren Forbes Beadle, Katherine M. Esposito, Christen K. Mirth, Coral G. Warr, Travis K. Johnson

## Abstract

Blood cells, known as hemocytes in invertebrates, play important and conserved roles in immunity, wound healing and tissue remodelling. The control of hemocyte number is therefore critical to ensure these functions are not compromised, and studies using *Drosophila* are proving useful for understanding how this occurs. Recently, the well characterised embryonic patterning gene, *torso-like* (*tsl*), was identified as being required both for normal hemocyte number and for providing immunity against certain pathogens. Here, we report that Tsl is required specifically during the larval phase of hematopoiesis, and that the reduced hemocyte number found in *tsl* mutant larvae is likely the result of a reduced larval growth rate and compromised insulin signalling. Consistent with this, we find that impairing insulin-mediated growth, either by nutrient deprivation or genetically, results in fewer hemocytes. This is likely the result of impaired insulin-like signalling in the hemocytes themselves since modulation of Insulin-like Receptor (InR) activity specifically in hemocytes causes concomitant changes to their population size in developing larvae. Taken together, our work reveals the strong relationship that exists between body size and hemocyte number, and suggests that insulin-like signalling contributes to, but is not solely responsible for, keeping these tightly aligned during larval development.

## Introduction

Blood cells perform a host of critical homeostatic and developmental functions. These include the phagocytosis of pathogens, cellular debris and other materials, and the secretion of histamine, cytokines and antimicrobial peptides to aid in immunity. In insects such as the fruit fly *Drosophila melanogaster*, these functions are performed by cells called hemocytes that circulate in the hemolymph both throughout development and in the adult. *Drosophila* have three types of mature hemocytes: plasmatocytes, crystal cells and lamellocytes. Plasmatocytes are phagocytic macrophage-like cells that make up >90% of the total hemocyte population, and function in pathogen elimination and tissue remodelling (Lanot et al. 2001, Tepass et al. 1994). Crystal cells facilitate wound healing and pathogen neutralisation via melanisation, and comprise <5% of hemocytes (Rämet et al. 2002, Lanot et al. 2001, Rizki, Rizki, and Grell 1980). Lamellocytes are large highly specialised hemocytes that function to encapsulate wasp eggs, however they typically only appear upon wasp parasitisation (Rizki and Rizki 1992).

All hemocytes in the adult fly derive from two populations of undifferentiated cells called prohemocytes that are specified at different stages of development (Holz et al. 2003). The first prohemocyte population originates in the procephalic mesoderm of the embryo (Tepass et al. 1994). Here, prohemocytes proliferate and differentiate into plasmatocytes and crystal cells, which then migrate throughout the embryo (Tepass et al. 1994). Embryonic plasmatocytes, but not crystal cells, persist into the larval stages and undergo a phase of proliferative expansion as the larva grows (Lebestky et al. 2000, Lanot et al. 2001, Holz et al. 2003). During this stage, these plasmatocytes can either circulate within the hemolymph or reside sessile in epidermal-muscular pockets at the larval body wall, where they are known to proliferate at an increased rate (Lanot et al. 2001, Makhijani et al. 2011). Crystal cells are also produced from these plasmatocytes in the larva via transdifferentiation (Leitao and Sucena 2015). The second population of prohemocytes is found in the larval lymph gland. Here, prohemocytes proliferate and differentiate into both crystal cells and plasmatocytes, which are then released into circulation during metamorphosis (Holz et al. 2003).

Hemocytes are necessary to maximise an individual’s ability to survive upon infection. For example, the ablation of post-embryonic hemocytes increases the proportion of flies that die upon infection by several different pathogens (Charroux and Royet 2009, Defaye et al. 2009). While much attention has been given to the functions of hemocytes in immunity, we know relatively little about the factors and mechanisms that control hemocyte population size. There are several signalling pathways are currently known to control *Drosophila* hemocyte number. These include the Platelet Derived Growth Factor/Vascular Endothelial Growth Factor Receptor (Pvr) pathway, which acts to maintain hemocyte survival during embryogenesis (Brückner et al. 2004), and the Activin-β pathway, which coordinates larval hemocyte proliferation by facilitating the accumulation of hemocytes in sessile pockets (Makhijani et al. 2017). Given the complexity and size of the cellular immune system, and its importance to individual fitness, it is likely that there are many factors and pathways still to be identified that serve to control hemocyte numbers.

Recently, *torso-like* (*tsl*) mutant larvae were shown to have a reduced number of circulating plasmatocytes and crystal cells, implicating Tsl as a novel regulator of hemocyte number (Forbes-Beadle et al. 2016). Tsl is a member of the Membrane Attack Complex/Perforin-like (MACPF) protein family, and has been shown to control a variety of developmental processes by modulating the activity of several different cell signalling pathways. These include terminal patterning in the early embryo via the Torso receptor (Stevens et al. 1990), embryonic morphogenesis via Fog signalling (Johnson et al. 2017), and larval growth via Insulin-like signalling (Johnson et al. 2013, Henstridge et al. 2018). Due to the latter role, *tsl* mutants have a reduced body size. We therefore wondered if the reduced hemocyte number observed in *tsl* mutant animals might be the result of the reduction in body size. Here, we show that this is indeed the case, and we further find that the same phenotype occurs upon modulating insulin-like signalling, both systemically and directly in hemocytes. Together, these data suggest that *tsl* acts with the insulin-like signalling pathway and that these contribute to coordinating hemocyte number and body size.

## Materials and methods

### Drosophila stocks and maintenance

The following stocks were from the Bloomington *Drosophila* Stock Centre: *w*^1118^ (BL3605), *hml-GAL4,UAS-GFP* (BL6397), *hmlΔ-GAL4,UAS-GFP* (BL30142), *UAS-RedStinger* (BL8546), *Df(3R)caki*^X-313^ (BL6784; called *tsl*^*caki*^ here), *chico*^*1*^ (BL10738), *Df(2L)BSC143* (BL9503; called *chico*^*df*^ here), UAS-InR^CA^ (BL8263) and UAS-InR^DN^ (BL8251). *srpHemo-GAL4* and *hmlΔ-dsRed* were kind gifts from Katja Brückner (Brückner et al. 2004, Makhijani et al. 2011). *tsl*^Δ^ is as previously described (Johnson et al. 2013). *chico*^*1*^ and *chico*^*df*^ were maintained over green balancers and experiments conducted on transheterozygous *chico*^*1*^/*chico*^*df*^ mutants as both *chico*^*1*^ and *chico*^*df*^ homozygotes were weak. *tsl*^Δ^, *tsl*^*caki*^ and *hmlΔ-GAL4,UAS-GFP* were maintained over TM6B. Flies were raised and maintained on standard sucrose, yeast and semolina fly media except where otherwise stated. For all experiments flies were maintained at 25°C except for the InR transgene expression experiment, which was performed at 29°C.

### Embryonic hemocyte imaging and quantification

Embryos laid by females from a cross between *srpHemo-Gal4*, which expresses in embryonic hemocytes (Brückner et al. 2004), and *UAS-RedStinger* lines were collected on apple juice agar supplemented with yeast paste overnight. Embryos were dechorionated in 50% vol/vol bleach before fixation in phosphate buffered saline (PBS) with 4% paraformaldehyde and an equal volume of heptane for 30 minutes. Vitelline membranes were removed by the addition of methanol before rehydration in PBS with 0.1% Triton X-100. Embryos were mounted in Vectashield and imaged at 20x magnification on an Olympus CV1000 confocal microscope. Maximum projection images were generated from 12 z-slices per embryo (stage 16) with at least 17 embryos sampled for each genotype. Hemocytes were counted using the ‘Image-based Tool for Counting Nuclei (ITCN)’ plugin for ImageJ (Schneider, Rasband, and Eliceiri 2012).

Embryonic hemocyte migration patterns were recorded by live-imaging embryos of the genotype described above following a two-hour egg collection. Embryos then developed for four hours before their dechorionation and mounting in an 8-well slide (Ibidi) containing PBS. Embryos were imaged for 9 hours (from approximately stage 10 to 16) at 20x magnification using a confocal microscope (CV1000). Twenty z-slices with a z-step distance <3.5μm were acquired at 5-minute intervals for each embryo and at least 15 embryos were imaged per genotype.

### Larval weighing and hemocyte quantification

Embryos were laid on apple juice agar supplemented with yeast paste for 24 hours. 0-24 hour old larvae were collected, selecting against green balancers where required, and reared in non-crowded, density-controlled conditions. Wandering stage larvae were used for larval hemocyte analyses except where stated otherwise, selecting against TM6B larvae where required. To weigh larvae, individuals were first washed in PBS, checked to ensure that all debris was removed, blotted dry and weighed on an ultra microbalance (XP2U, Mettler Toledo) immediately prior to bleeding.

Fluorescent circulating and sessile hemocytes were quantified following their extraction from larvae containing either *hml-GAL4,UAS-GFP; hmlΔ-GAL4,UAS-GFP;* or *hmlΔ-dsRed* transgenes, which all express in larval hemocytes (Goto, Kadowaki, and Kitagawa 2003, Sinenko and Mathey-Prevot 2004, Makhijani et al. 2011). To extract circulating hemocytes, larvae were washed once in PBS, transferred individually to an 8-well slide (Ibidi) containing PBS, and then bled for at least two minutes through a hole torn in their dorsal-posterior cuticle. Sessile hemocytes were then removed by scraping the remaining hemocytes from the carcass into a new well containing PBS, being careful to avoid disruption to lymph glands, as previously described (Petraki, Alexander, and Brückner 2015). Hemocytes were resuspended by pipetting to minimise clumping, allowed to settle for 10 minutes, then the entire well was imaged using a Leica AF6000 LX for the nutrient deprivation and InR transgene expression experiments, or an Olympus CV1000 for all other experiments. The number of fluorescent cells per well was quantified using ImageJ as previously described (Petraki, Alexander, and Brückner 2015). The total number of hemocytes per larva was calculated as the sum of the circulating and sessile hemocytes.

## Results

We previously reported that *tsl* mutant adults have compromised immunity and a reduced ability to clear pathogens, likely caused by the markedly reduced number of circulating larval plasmatocytes and crystal cells found in these animals (Forbes-Beadle et al. 2016). Since circulating larval hemocytes originate from the embryonic rather than lymph gland hemocyte lineage, Tsl must control circulating hemocyte number by acting either during the embryonic or larval stages of hematopoiesis. To distinguish between these possibilities, we first measured the hemocyte number in *tsl*^*Δ*^ homozygous mutant embryos while marking their hemocytes using *srpHemo-GAL4*>*UAS-RedStinger*. We found no difference in hemocyte number between *tsl*^*Δ*^*/tsl*^*Δ*^ and control *tsl*^*Δ*^/+ embryos (Fig. 1A, p = 0.821). We also used live-imaging to track hemocyte migration patterns and found no migration defects in *tsl*^*Δ*^ *tsl*^*Δ*^ embryos compared to control *tsl*^*Δ*^/+ embryos (Fig. 1B). These data suggest that Tsl does not influence embryonic hemocyte development, and therefore may act during larval hematopoiesis.

**Figure 1.**
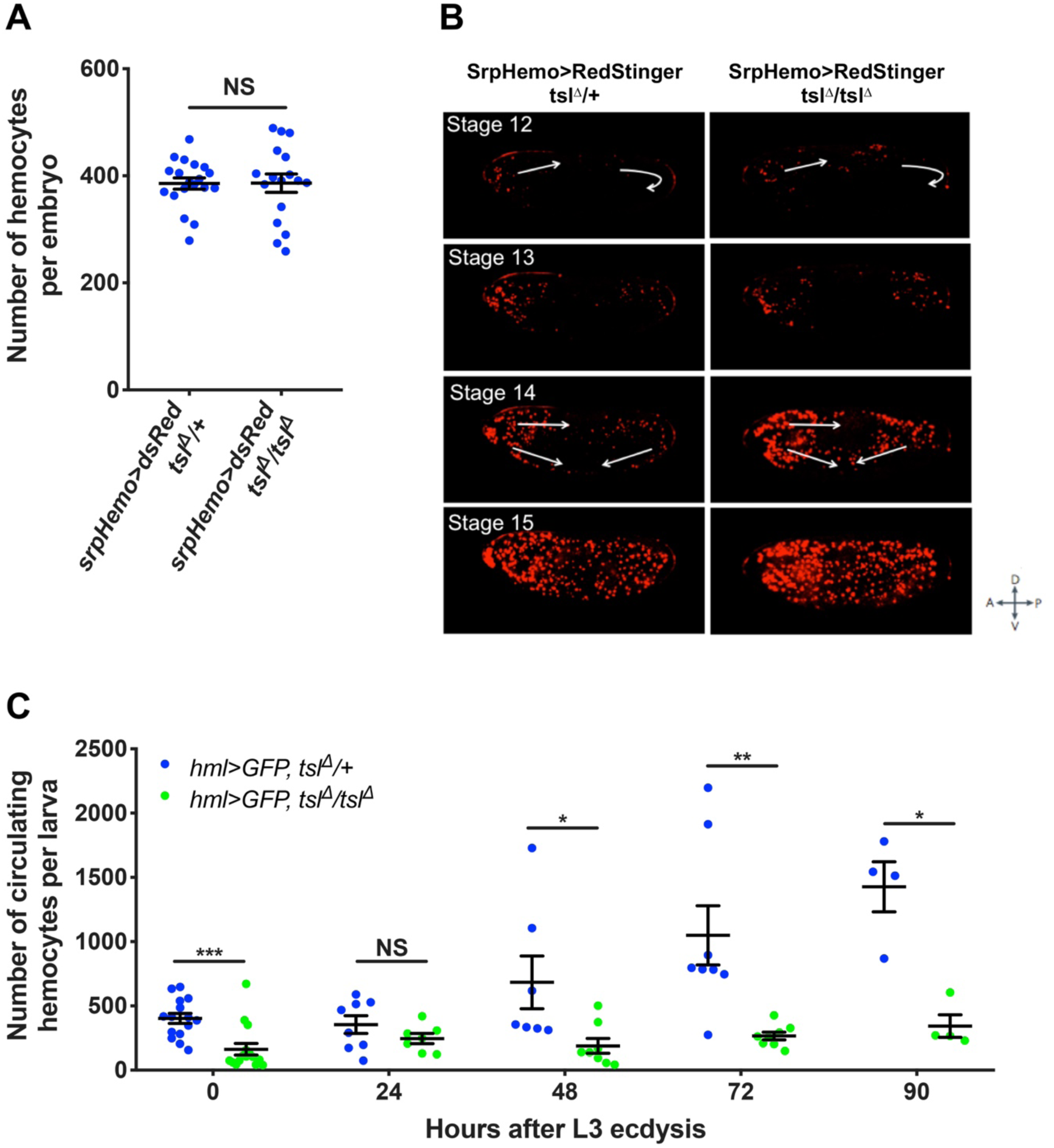
Tsl influences larval hemocyte numbers. (**A**) Hemocyte numbers in stage 16 *tsl*^*Δ*^*/tsl*^*Δ*^ and *tsl*^*Δ*^/+ embryos (*srpHemo-GAL4*>*UAS-RedStinger*). No difference in hemocyte number was observed. (p=0.821, n≥17). (**B**) Stills from live-imaging of stage 12-15 *tsl*^*Δ*^*/tsl*^*Δ*^ and *tsl*^*Δ*^/+ embryos. Arrows indicate the movement of hemocytes. No difference in migration pattern was observed between *tsl*^*Δ*^*/tsl*^*Δ*^ and *tsl*^*Δ*^/+ control embryos. (**C**) Circulating hemocyte counts from *tsl*^*Δ*^*/tsl*^*Δ*^ and *tsl*^*Δ*^/+ larvae at regular time points after transitioning to the third larval stage (L3) of development (n≥4). NS, not significant, *p<0.05, **p<0.01, ***p<0.001, Mann-Whitney test. Data are mean ±1 standard error.

To test this, we next counted circulating hemocytes in *tsl*^*Δ*^*/tsl*^*Δ*^ larvae throughout the third instar larval stage (using *hemolectin (hml)-GAL4*>*UAS-GFP*). A significant reduction in circulating hemocyte numbers was apparent in *tsl*^*Δ*^*/tsl*^*Δ*^ larvae upon moulting to the third instar stage compared to *tsl*^*Δ*^/+ controls (Fig. 1C, p<0.001). This difference became more pronounced in the late third instar stage (Fig. 1C, 72h, p=0.001), suggesting that Tsl may be continually required for the expansion of the hemocyte population throughout late larval development.

Recent work has shown that the larval hemocytes that reside in sessile patches proliferate more rapidly than those in circulation, and an inability of hemocytes to associate in sessile patches can disrupt their relative numbers and subsequently overall larval hemocyte number (Makhijani et al. 2011, Makhijani et al. 2017). This prompted us to explore whether *tsl* influences the sessile hemocyte population (*hmlΔ-dsRed*). We found that *tsl*^*Δ*^*/tsl*^*Δ*^ larvae indeed have a greatly reduced number of sessile hemocytes compared to *tsl*^*Δ*^/+ controls (Fig. 2A, p=0.003). We also asked if Tsl affects the accumulation of hemocytes in sessile patches by determining the proportion of hemocytes in circulation relative to the total number. This showed that the proportion of total hemocytes that are in circulation in *tsl*^*Δ*^*/tsl*^*Δ*^ larvae is not different to that observed in *tsl*^*Δ*^/+ control larvae (Fig. 2B, p=0.573). Thus, *tsl* is required for the expansion of both the circulating and sessile hemocyte populations, and is unlikely to influence hemocyte accumulation in sessile patches.

**Figure 2.**
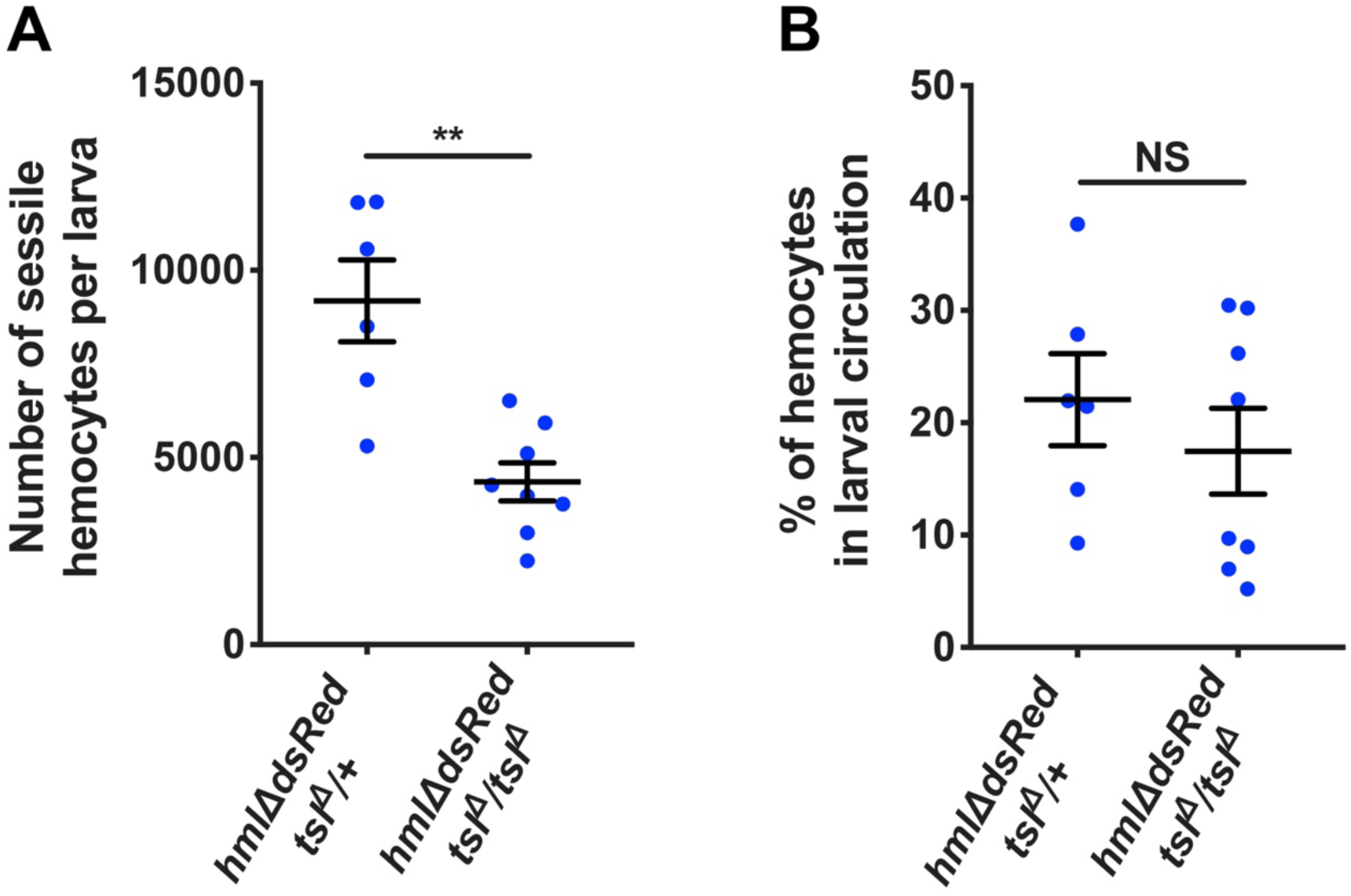
Tsl is required for both sessile and circulating larval hemocyte populations. (**A**) Quantification of sessile hemocyte numbers (*hmlΔ-dsRed*) in *tsl*^*Δ*^*/tsl*^*Δ*^ and *tsl*^*Δ*^/+ larvae. *tsl*^*Δ*^*/tsl*^*Δ*^ larvae have significantly fewer sessile hemocytes than *tsl*^*Δ*^/+ controls (p=0.003). (**B**) No significant difference in the proportion of hemocytes in circulation was observed between *tsl*^*Δ*^*/tsl*^*Δ*^ mutant and *tsl*^*Δ*^/+ control larvae (p=0.573). NS, not significant, *p<0.05, **p<0.01, ***p<0.001, Mann-Whitney test. Data are mean ±1 standard error (n≥6).

*tsl*^*Δ*^ mutant animals have previously been reported to have a reduced growth rate and body size due to a disruption in the insulin-like signalling pathway (Henstridge et al. 2018, Johnson et al. 2013). Therefore, we questioned whether this could be contributing to the larval hemocyte number deficit. Since both weight and hemocyte number are quantitative traits we tested this by weighing individual *tsl*^*Δ*^/+ and *tsl*^*Δ*^*/tsl*^*Δ*^ larvae (as a proxy for body size and therefore growth) and counting their total hemocyte number (i.e. circulating and sessile, *hmlΔ-dsRed*). We found that hemocyte number increases linearly with body size (Fig. 3A, log(weight) effect p=0.004, Table S1) and this relationship is the same for both genotypes (Fig. 3A; Table S1, weight*genotype effect p=0.594). However, for any given weight, *tsl*^*Δ*^*/tsl*^*Δ*^ larvae have fewer hemocytes when compared to *tsl*^*Δ*^/+ controls (Fig. 3A; Table S1, genotype effect p<0.001). To determine if this weight-independent effect was due specifically to loss of *tsl* or other elements in the genetic background, we repeated the test using a transheterozygous *tsl* allele combination (*tsl*^*Δ*^/*tsl*^*caki*^; *hmlΔ-dsRed*). This confirmed that these larvae also have a reduction in total hemocyte number compared to both heterozygote controls (Fig. 3B, p<0.001), and the effect of weight on hemocyte number was again observed (Fig. 3C; Table S2, log(weight) effect p=0.034). However, for this genotype we did not detect a weight-independent effect of *tsl* on hemocyte number (Fig. 3C; Table S2, genotype effect p=0.007 and Tukey post-hoc p>0.05 for all genotypic combinations). This suggests that there may be other genetic changes present in the *tsl*^*Δ*^*/tsl*^*Δ*^ background that influence hemocyte number independently of an effect on body size. Thus, overall these data strongly suggest that *tsl* influences hemocyte number via its effect on larval body size.

**Figure 3.**
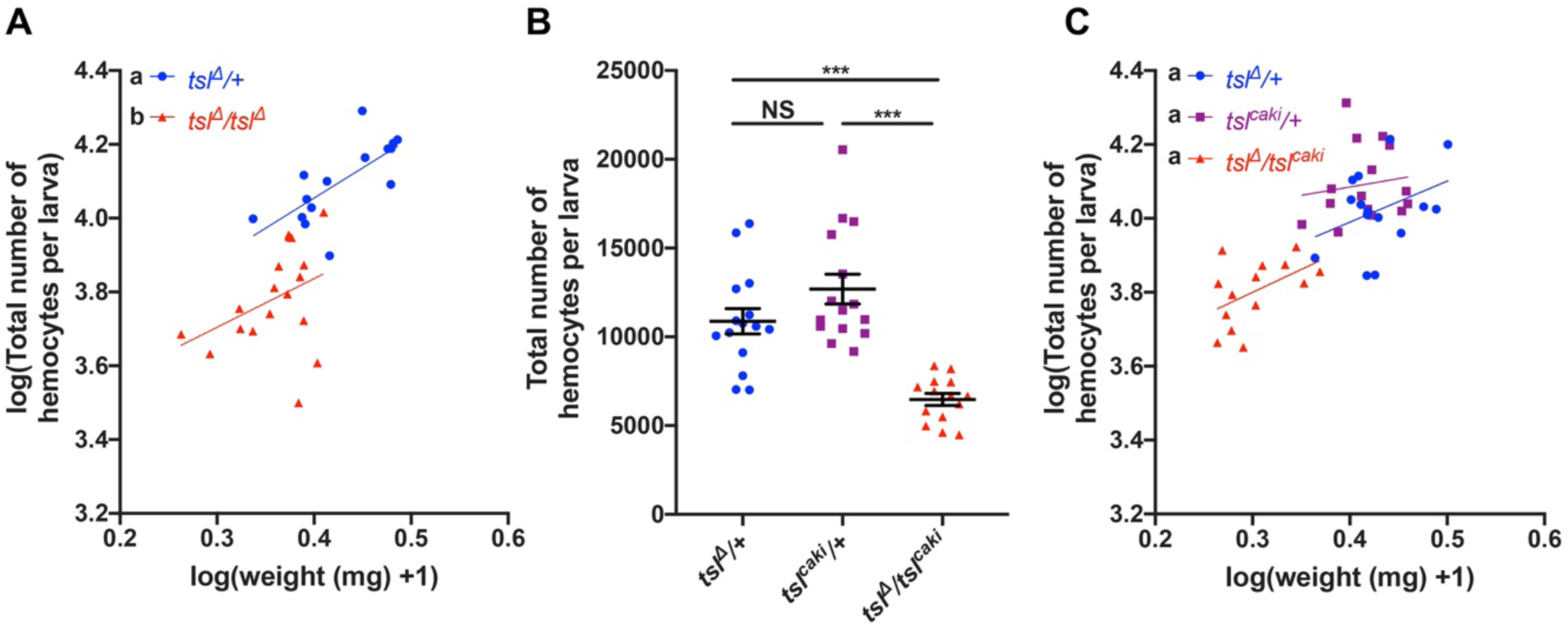
The reduced hemocyte number in *tsl* mutants is the result of larval growth impairment. (**A**) Quantification of weight and total hemocyte number (*hmlΔ-dsRed*) in individual *tsl*^*Δ*^*/tsl*^*Δ*^ and *tsl*^*Δ*^/+ larvae. While a relationship exists between total hemocyte number and weight, for any given weight *tsl*^*Δ*^*/tsl*^*Δ*^ mutant larvae have fewer hemocytes than their heterozygous counterparts. (**B**) Analysis of *tsl*^*Δ*^/+, *tsl*^*caki*^/+ and *tsl*^*Δ*^/*tsl*^*caki*^ larvae (*hmlΔ-dsRed*) reveals that *tsl*^*Δ*^/*tsl*^*caki*^ larvae have significantly fewer hemocytes than both heterozygous controls (p<0.001, n≥14). (**C**) Incorporating larval weight into this analysis revealed that *tsl*^*Δ*^/*tsl*^*caki*^ larvae did not have fewer hemocytes than their heterozygous counterparts at any given weight. For (**A**) and (**C**) regression lines that cannot be represented by the same line of best fit are indicated by different letters. For (**B**), NS, not significant, *p<0.05, **p<0.01, ***p<0.001, Mann-Whitney test. Data are mean ±1 standard error.

Since *tsl* has been shown to mediate growth and final body size via the insulin-like signalling pathway, we next investigated whether insulin-like signalling also impacts the larval hemocyte population. For this we quantified weight and hemocyte numbers in larvae that were mutant for *chico*, which encodes an adaptor protein required for transduction of the InR signal (Böhni et al. 1999). Consistent with previous reports for *chico* mutant animals (Böhni et al. 1999), we found that transheterozygous *chico* mutants (*chico*^*1*^*/chico*^*def*^) are significantly smaller than both controls (Fig. 4A; p<0.001). In addition, *chico*^*1*^*/chico*^*def*^ larvae exhibited a significant reduction in hemocyte number compared to both heterozygote controls (*hmlΔ-GAL4*>*UAS-GFP*, Fig. 4B; p<0.001). This effect is explained by their reduced size, with no weight-independent effects observed (Fig. 4C; Table S3, log(weight) effect p<0.001, genotype effect p=0.322). Thus, both loss of *tsl* and reduced insulin-like signalling impacts the size of the hemocyte population via their effect on whole animal body size.

**Figure 4.**
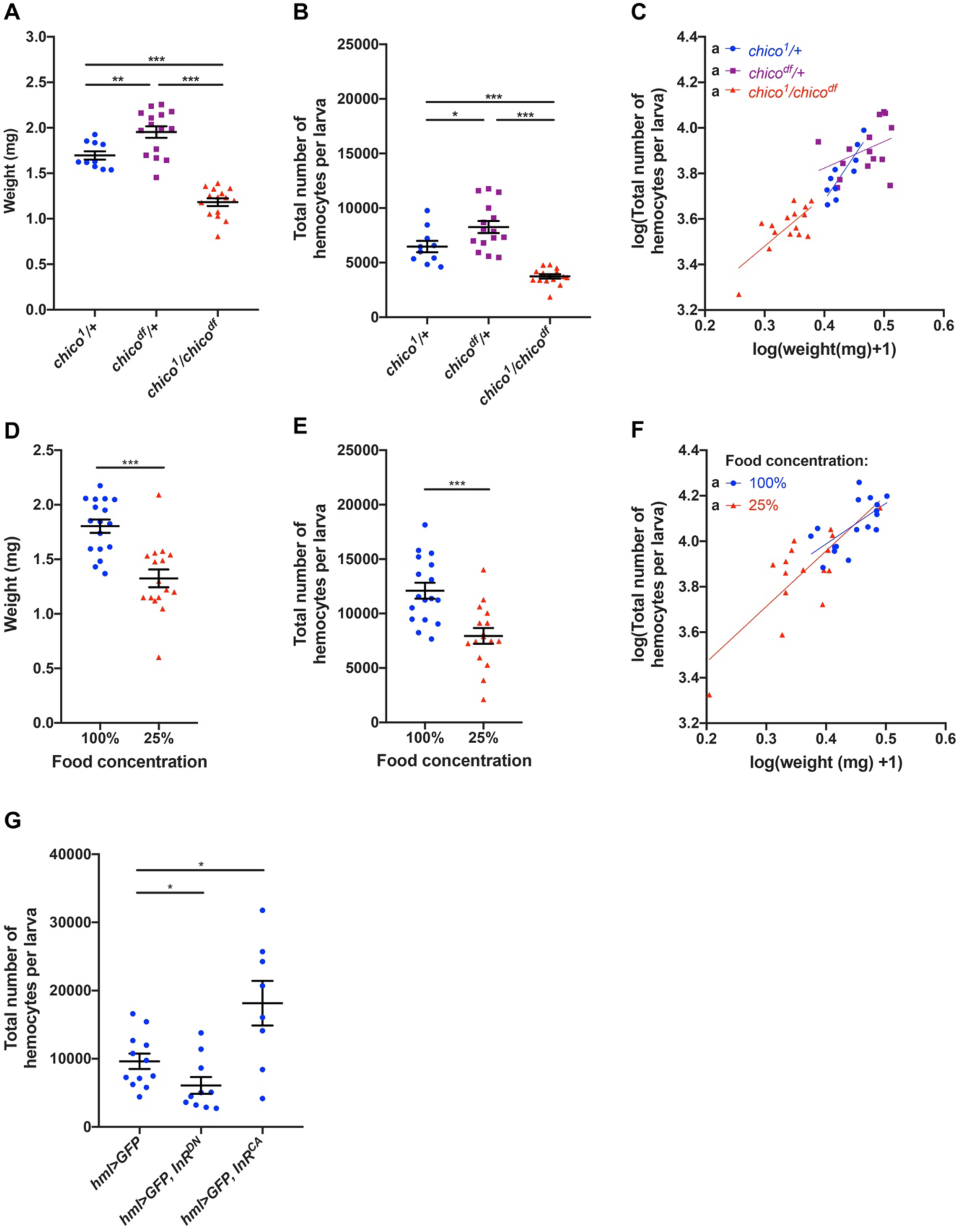
Hemocyte-autonomous insulin-like signalling links larval growth to hemocyte number. (**A**) *chico*^*1*^*/chico*^*def*^ larvae are significantly smaller than both heterozygous controls (p<0.001, n≥10) (**B**) *chico*^*1*^*/chico*^*def*^ larvae have significant fewer hemocytes than both heterozygous controls (*hmlΔ-GAL4*>*UAS-GFP*, p<0.001, n≥10). (**C**) Incorporating larval weight into this analysis reveals that *chico*^*1*^*/chico*^*def*^ larvae did not have fewer hemocytes than their heterozygous counterparts at any given weight. (**D**) Larvae raised on a diluted diet (25%) were significantly smaller than larvae raised on the standard 100% diet (p<0.001, n≥16). (**E**) Larvae raised on a 25% diet had significantly fewer hemocytes than fully fed controls (*hmlΔ-dsRed*, p<0.001). **(F**) Incorporating larval weight into this analysis reveals that larvae raised on the 25% diet did not have fewer hemocytes than those on the 100% diet at any given weight. (**G**) Quantification of total hemocyte number (*hml-GAL4*>*UAS-GFP*) in larvae expressing either a dominant negative or constitutively active form of InR (*InR*^*DN*^ and *InR*^*CA*^, respectively) in hemocytes. Expression of *InR*^*DN*^ resulted in a reduction in total hemocyte number (p=0.030), while expression of *InR*^*CA*^ resulted in an increase (p=0.039, n≥8). For (**B**) and (**E**) regression lines that cannot be represented by the same line of best fit are indicated by different letters. For (**A**), (**C**), (**D**) and (**F**), NS, not significant, *p<0.05, **p<0.01, ***p<0.001, Mann-Whitney test. Data are mean ±1 standard error.

Nutrient deprivation also causes a reduction in larval weight and body size due to altered levels of insulin-like signalling (Hietakangas and Cohen 2009). Therefore, as an alternative means of inducing insulin-dependent growth impairment, we next quantified hemocytes bled from wildtype larvae (*hmlΔ-dsRed*) fed on a nutrient deprived diet compared to a normal diet. As expected, we found that reducing the concentration of our standard diet to 25% of normal nutrient levels significantly reduced larval weights (Fig. 4D; p<0.001). We observed corresponding significant reductions in hemocyte number in these larvae (Fig. 4E; p<0.001). The variation in hemocyte number was largely attributed to weight regardless of the diet on which the larva was raised (Fig. 4F; Table S4, log(weight) effect p<0.001, diet effect p=0.663). Together, these data further support the idea that insulin signalling links the size of the hemocyte population to overall larval growth.

We next asked if insulin signalling was regulating hemocyte population size via a cell autonomous role in the hemocytes themselves, or indirectly via roles in other tissues. To test this we expressed transgenes carrying dominant negative and constitutively active forms of InR (InR^DN^ and InR^CA^, respectively) specifically in larval hemocytes using *hml-GAL4*. Reducing insulin-like signalling by expression of InR^DN^ resulted in a significant reduction in total larval hemocyte number (*hml-GAL4*>*UAS-GFP*, Fig. 4G; p=0.030), while overactivation of insulin-like signalling via expression of InR^CA^ had the opposite effect (Fig. 4G; p=0.039). These data strongly suggest that, during the larval stages, hemocyte-autonomous insulin-like signalling is important for the regulation of their overall hemocyte numbers.

## Discussion

Here we provide the first evidence for a strong relationship between body size and hemocyte number in *Drosophila* larvae. Firstly, this highlights the importance of accounting for differences in larval weight when using whole animal mutant lines to identify novel regulators of larval hemocyte number. We have found that up to 80% of the variation observed in larval hemocyte number within genotypes is due solely to variation in larval weight, despite controlling for factors that influence body size variation such as larval culture density (Santos, Fowler, and Partridge 1994). We therefore recommend using an approach that accounts for body size variation to enable more subtle changes to larval hemocyte number to be detected.

Secondly, we find that the relationship between body size and hemocyte number is linear, suggesting that it is important for larvae to keep hemocyte concentration (hemocytes per unit weight) constant no matter the body weight attained. This is consistent with what is observed in mammals. For example, leukocyte concentration remains constant regardless of body size in humans (Chmielewski et al. 2017). What drives organisms to have an optimal hemocyte/blood cell concentration? In addition to their role in immunity, hemocytes have numerous other duties (e.g. extracellular matrix deposition), and thus too many or too few hemocytes may result in both immunity and non-immunity related homeostatic imbalances (Banerjee et al. 2019). For example, since hemocytes (and leukocytes more generally) are known to be high energy users, it may be metabolically detrimental to have too many hemocytes (Dolezal et al. 2019). In support of this, a recent study showed that increases in hemocyte number cause sensitivity to nutrient deprivation (Ramond et al. 2020). Thus, the determinants of the optimal hemocyte concentration may be, at least in part, the result of competing energy needs between to roles of hemocytes and the growth of tissues.

Finally, we show that both *tsl* and insulin-like signalling influence hemocyte numbers via their effect on larval growth. Insulin-like signalling plays an important role in the coordination of growth for many organs in response to changes in extrinsic factors such as nutrient availability (Hietakangas and Cohen 2009). Our work suggests that the circulating and sessile hemocyte populations in larvae may be coordinated similarly. This is perhaps unsurprising given that the insulin-like peptides responsible for mediating a large proportion of InR-dependent growth are in circulation and therefore in direct contact with hemocytes (Rulifson, Kim, and Nusse 2002). Interestingly however, our data also revealed that hemocyte number and body size remain coordinated even when insulin-like signalling is compromised. This strongly suggests that additional growth pathways are likely to be involved in maintaining normal hemocyte concentrations. One recently identified factor that may play a role here is NimB5, which is secreted from fat body cells in a nutrient-dependent manner to dampen hemocyte proliferation and avoid energy wastage (Ramond et al 2020). It will be interesting to learn how NimB5, insulin-like signalling and other pathways control hemocyte numbers with respect to *Drosophila* growth and determine the extent to which they are conserved across animals.

## Abbreviations

Tsl: Torso-like
InR: Insulin-like receptor
hml: hemolectin
Pvr: Platelet Derived Growth Factor/ Vascular Endothelial Growth Factor Receptor

## Acknowledgments

We would like to thank the Australian *Drosophila* Biomedical Research Facility (OzDros) and Monash Micro Imaging for technical support. In addition, we thank the Bloomington *Drosophila* Stock Centre and Katja Brückner for fly stocks, and Grace Jefferies for critical reading of the manuscript. This work was supported by the Monash University Science-Medicine, Nursing, and Health Science Faculties Interdisciplinary Research Scheme.

