## Supplemental data for "Insulin-like signalling influences the coordination of *Drosophila* larval hemocyte number with body size"

**Table S1**

| $tsl^{\Delta}$ vs $tsl^{\Delta}$/+ | | | |
|---|---|---|---|
| | **Sum of squares** | **F value** | **Pr(>F)** |
| **$\log_{10}$(Weight)** | 0.638 | 9.695 | 0.004 ** |
| **Genotype** | 1.261 | 19.152 | <0.001 *** |
| **$\log_{10}$(Weight)*Genotype** | 0.019 | 0.291 | 0.594 NS |

Data was fit with the linear model $\log_{10}$(total hemocyte number) = $\log_{10}$(weight) + genotype + $\log_{10}$(weight)*genotype ($F_{3, 29}$ = 25.55, Adjusted $R^2$ = 0.70) using RStudio.

**Table S2**

| ***tsl**$^{caki}$***/+ vs *tsl**$^{\Delta}$*/+ vs *tsl**$^{\Delta}$*/tsl**$^{caki}$* | | | |
|---|---|---|---|
| | **Sum of squares** | **F value** | **Pr(>F)** |
| **log$_{10}$(Weight)** | 0.246 | 4.841 | 0.034 * |
| **Genotype** | 0.568 | 5.584 | 0.007 ** |
| **log$_{10}$(Weight)*Genotype** | 0.021 | 0.203 | 0.817 NS |
| **Genotype Tukey post-hoc** | | | |
| | | **T ratio** | **P value** |
| ***tsl**$^{caki}$*/+ vs *tsl**$^{\Delta}$*/+** | | 1.994 | 0.128 NS |
| ***tsl**$^{caki}$*/+ vs *tsl**$^{\Delta}$*/tsl**$^{caki}$*** | | 2.994 | 0.069 NS |
| ***tsl**$^{\Delta}$*/+ vs *tsl**$^{\Delta}$*/tsl**$^{caki}$*** | | 0.914 | 0.635 NS |

Data was fit with the linear model log$_{10}$(total hemocyte number) = log$_{10}$(weight) + genotype + log$_{10}$(weight)*genotype ($F_{5, 38}$ = 14.67, Adjusted $R^2$ = 0.61) using RStudio. This was followed with a Tukey posthoc test to determine which genotype(s) had significantly different effects on total hemocyte number compared to the others.

**Table S3**

| ***chico*$^{1}$*/+* vs *+/chico*$^{def}$ vs *chico*$^{1}$*/chico*$^{def}$** | | | |
|---|---|---|---|
| | **Sum of squares** | **F value** | **Pr(>F)** |
| **log$_{10}$(Weight)** | 0.806 | 22.041 | <0.001*** |
| **Genotype** | 0.086 | 1.173 | 0.322 NS |
| **log$_{10}$(Weight)*Genotype** | 0.152 | 2.076 | 0.141 NS |

Data was fit with the linear model log$_{10}$(total hemocyte number) = log$_{10}$(weight) + genotype + log$_{10}$(weight)*genotype ($F_{5, 34}$ = 31.38, Adjusted $R^2$ = 0.80) using RStudio.

**Table S4**

| **100% diet vs 25% diet** | | | |
|---|---|---|---|
| | **Sum of squares** | **F value** | **Pr(>F)** |
| **log$_{10}$(Weight)** | 2.362 | 40.944 | <0.001*** |
| **Diet** | 0.011 | 0.194 | 0.663 NS |
| **log$_{10}$(Weight)*Diet** | 0.011 | 0.192 | 0.665 NS |

Data was fit with the linear model log$_{10}$(total hemocyte number) = log$_{10}$(weight) + diet + log$_{10}$(weight)*diet ($F_{3, 29}$ = 24.31, Adjusted $R^2$ = 0.69) using RStudio.
